# Functional synapse types via characterization of short-term synaptic plasticity

**DOI:** 10.1101/648725

**Authors:** Jung Hoon Lee, Luke Campagnola, Stephanie C. Seeman, Tim Jarsky, Stefan Mihalas

## Abstract

The strengths of synaptic connections dynamically change depending on the history of synaptic events, which is referred to as short-term plasticity (STP). While STP’s underlying mechanisms are well researched, its exact functions remain poorly understood. This is in part due to the diverse patterns of STP experimentally reported. Recently, the Allen Institute for Brain Science has launched the synaptic physiology pipeline to characterize the diverse properties of synapses. Since this pipeline generates a large-scale survey of synapses in mouse primary visual cortex using highly standardized experimental protocols, it provides a unique opportunity to study diverse patterns of STP. Here, we develop an end-to-end workflow that can characterize STP from the Allen Institute for Brain Science pipeline data and conduct network simulations to infer STP’s functions. Employing this workflow, we find 1) that diverse patterns of STP exist even in the same synapse classes and 2) that postsynaptic neurons’ responses have distinct characteristics depending on STP.

## 1 Introduction

Synapses mediate spikes from presynaptic neurons to postsynaptic neurons by consuming synaptic resources. As the available synaptic resources depend on the history of synaptic events, the efficacies of synapses (synaptic inputs) change dynamically. This resource-related change is restored very rapidly (usually within seconds) and referred to as short-term plasticity (STP). Due to its fast recovery to baseline states, STP cannot support learning in the brain, as long-term plasticity does. Instead, STP has been thought to perform gain control [1] and serve temporary memory buffer [14], but its functions are poorly understood in part because the characterization of its diverse patterns remain incomplete.

Multiple studies[1, 10, 24, 26] have proposed mathematical models to account for changes in synaptic event strength, raising the possibility of mathematical characterization of STP. Among them, the phenomenological model (TM model) proposed by Tsodyks et al. [26] has been considered the most successful one and served as a standard model of STP. However, even the TM model cannot explain all experimental observations [6, 11, 28]; see also [10]. For instance, the recovery of depletion is faster when presynaptic neurons are in a high firing regime [6, 25], and the TM model cannot address this effective facilitation occurring selectively. Yet, mathematical models can address such newly observed dynamics [8, 10]. That is, it is possible, in principle, to mathematically characterize diverse patterns of STP. However, it should be noted that mathematical modeling alone is not enough for proper characterization of STP because it requires well-defined data.

Unfortunately, synaptic properties are difficult to measure due to the high variability and diversity of synaptic transmission. Then, how should we address this challenge? An earlier study [23] from the Allen Institute for Brain Science proposed a large-scale survey of synapses as an effective solution, which was collected using industrial-level standardized experimental protocols. Specifically, it showed 1) that a large-scale survey can help us identify synapse classes and their functions effectively and 2) that a mathematical model can be used to characterize STP classes. However, we note that the mathematical characterization of STP in this study [23] has limitations. First, it assumed that all synapses in synapse classes are homogeneous and did not address the possibility of heterogeneous synapses in the classes. Second, it utilized only one synapse model for all classes and consequently ignored the possibility that synapse classes can have independent synaptic dynamics.

Here, we address the limitations mentioned above by examining the homogeneity of synapse classes and use more comprehensive models of STP. Our new analyses suggest that synapse classes can be split into subclasses with distinct STP patterns. Further, to gain insights into functions of diverse STP, we construct a network model and simulate its responses. We believe that this end-to-end workflow will help us better understand STP’s contribution to neural dynamics and our cognitive functions in the end.

## 2 Methods

We start this section with a summary of experimental protocols; see [23] for details. In the original study [23], synaptic connections between excitatory neurons in layers (L) 2/3, 4 and 5 were probed to characterize STP. With a pair of excitatory neurons in the same class identified, postsynaptic potential (PSP) amplitudes were measured while 12 pulses were used to induce presynaptic neurons to generate spikes. A set of 12 pulses was referred to as a ‘sweep’, which we will also use in this study. In each sweep, 12 pulses were injected to presynaptic neurons at a fixed frequency (chosen from 10, 20, 50, 100 Hz). The delay periods were inserted between 8th and 9th pulses to estimate the speed of synapses’ recovery to their baseline states. In the sweeps with 50 Hz pulses, 5 delay periods (250, 500, 1000, 2000, 4000 ms) were used. In all other sweeps, the delay period was fixed at 250 ms. For each frequency of presynaptic pulses (i.e., stimulation protocol), PSPs were measured in at least 5 sweeps and averaged over them. Finally, PSPs were normalized to the PSP evoked by the first pulse to characterize STP; that is, the first PSP in each stimulation protocol is always fixed to be 1.

### 2.1 12 models of STP

According to the excellent review [10], at least three additional mechanisms need to be considered to explain STP further, and thus we construct synapse models with 5 synaptic dynamics (or alternatively, gating processes). Among them, the two dynamics are abstract versions of TM model, which were used in the earlier models [14, 20], and the rest are use-dependent replenishment, desensitization of receptors and slow modulation of release probability. These five temporal dynamics are shown in Eq. 1-5, respectively.

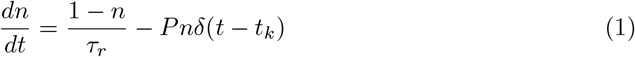

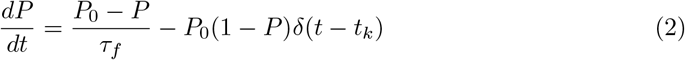

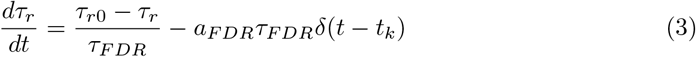

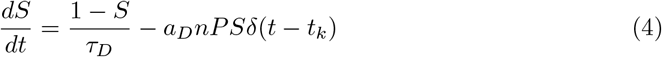

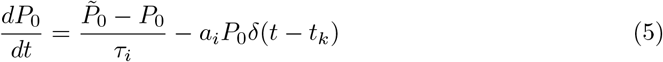

By choosing selectively from 5 gating processes, we build 12 different synapse models; the definition of 12 models are listed in Table 1.

**Table 1:** The definitions of synapse models. Each model includes a combination of 5 temporal dynamics, which are depression (DEP, Eq. 1), facilitation (FAC, Eq. 2), use-dependent replenishment (USR, Eq. 3), desensitization (DSR, Eq. 4) and slow modulation of release probability (SMR. Eq. 5).

| Model number | Temporal Dynamics | Model number | Temporal Dynamics |
| --- | --- | --- | --- |
| 1 | DEP | 7 | DEP, FAC, DSR |
| 2 | DEP, FAC | 8 | DEP, FAC, USR, DSR |
| 3 | DEP, USR | 9 | DEP, FAC, SMR |
| 4 | DEP, DSR | 10 | DEP, FAC, USR, SMR |
| 5 | DEP, USR, DSR | 11 | DEP, FAC, DSR, SMR |
| 6 | DEP, FAC, USR | 12 | DEP, FAC, USR, DSR, SMR |

### 2.2 X-means clustering

Unlike other conventional unsupervised clustering algorithms, which require a predefined number of clusters, ‘x-means’ clustering algorithm determines the optimal number of clusters [18]. In principle, it conducts k-means clustering and estimates Bayesian information criterion (BIC) for a wide range of potential numbers of clusters. The number of clusters, with which BIC is minimized, is selected as the optimal number of clusters. Following this idea, we conduct k-means clustering and calculate BIC depending on the number of clusters. In this study, we use scikit-learn [17] for k-means clustering and adopt BIC routine which is publicly available [15]. To minimize the stochastic bias from k-means clustering, we estimate BIC based on 1000 independent clustering results.

## 3 Results

To identify the functional synapse types (or groups), we re-analyze dynamics of synaptic events in mouse visual cortex published earlier [23]. The summary of the experimental protocols is given in Methods. We follow the original study to estimate normalized PSPs, and a set of normalized 12 PSPs collected in a sweep will be referred to as ‘PSP vector’ for the sake of brevity. That is, for a synapse, there are 5 PSP vectors collected with 50 Hz presynaptic spikes due to 5 different delay periods [23] and 1 PSP vector collected with all other presynaptic spikes. Below, we describe our results regarding subclasses of synapses, model-based characterization of STP and network models’ simulation conducted to gain insights into functions of STP.

### 3.1 Identifying homogeneous subclasses within synapse classes

In the original study [23], the connections between the identical excitatory neuron classes (types) were probed. Since STP is known to depend on pre and postsynaptic neurons [3, 9, 13, 19], synapse classes defined with presynaptic neuron class were used to characterize STP using these synapse classes. Specifically, there were four synapse classes 1) between excitatory neurons in L2/3, 2) between rorb-expressing neurons in L4, 3) between sim1-expressing neurons in L5, 4) and between tlx3-expressing neurons in L5. By characterizing STP of these classes, rather STP of individual synapses, it is possible to effectively address the stochasticity and diverse patterns of synaptic transmissions, if synapses in these classes are homogeneous. However, they [23] noted that the synapses between L2/3 neurons are poorly explained by the model, leading to the possibility that synapse classes defined by presynaptic neuron classes include heterogeneous subclasses.

To seek more precise characterizations of STP, we first ask if the four synapse classes (Syn_L23, Syn_rorb, Syn_sim1, Syn_tlx3) have heterogeneous subclasses. If synapses in the same class are homogeneous, the time courses of PSPs (i.e., PSP vectors) recorded using the same stimulation protocol would be identical throughout all synapses in the same class. Otherwise, the PSP vectors would be different from one another. With this possibility in mind, we turn to unsupervised clustering algorithms to test the homogeneity of synapses (i.e., the pattern of PSP changes). We note two potential issues in clustering synapses using PSP vectors. First, PSP vectors need to be collected with the same stimulation protocol. Otherwise, even PSP vectors from the same synapses will be different from each other, and clustering algorithms will consider them to belong to different synapses. That is, we need to select a single stimulation protocol for clustering. In this study, we use the first 8 components of PSP vectors collected with 50 Hz stimulation protocol to identify subclasses because there are 5 PSP vectors in the 50 Hz sweeps, and the first 8 components are collected under the identical stimulation condition, regardless of lengths of delay periods. Second, unsupervised learning algorithms require a predefined number of clusters (i.e., subclasses), but the actual numbers of subclasses are unknown. Since Pelleg and Moore [18] proposed to use BIC to find the optimal number of clusters for a given dataset, which was coined as the ‘x-means clustering’ (Methods), we use x-means clustering algorithms in this study. In principle, x-means employs k-means over a wide range of numbers of clusters to select the optimal number of clusters with BIC. That is, when the numbers of clusters are fixed, x-means is identical to k-means clustering. To remove the bias from the stochasticity of k-means clustering, we repeat x-means algorithms 1000 times and estimate the mean values of BICs from these 1000 trials.

As principal component analysis is commonly used for better clustering, we use 3 to 7 principal components to reduce the dimension of PSP vectors. Fig. 1 shows BIC values depending on the number of clusters and principal components for each synapse class. Using these values, we consider the 3 clusters and 3 principal components to be optimal for Syn_L2/3. We also consider 4 clusters and 7 principal components to be optimal for all other synapse classes (Syn_rorb, Syn_sim1, Syn_thx3). With these optimal numbers of clusters and principal components, we again conduct k-means clustering 3000 times for each synapse class and select the best clustering result from them using silhouette score. Many clustering results are identical in terms of silhouette score (Supplemental data), and we choose one randomly from the equally good results. During clustering, all PSP vectors are treated and clustered independently. Consequently, even PSP vectors from the same synapses can be assigned to multiple clusters (i.e., subclasses). We inspect how robustly PSP vectors from the same synapses are assigned to the same cluster and include synapses for further analysis only when 75 % of PSP vectors from the same synapses or more are assigned to the same cluster.

**Figure 1:**
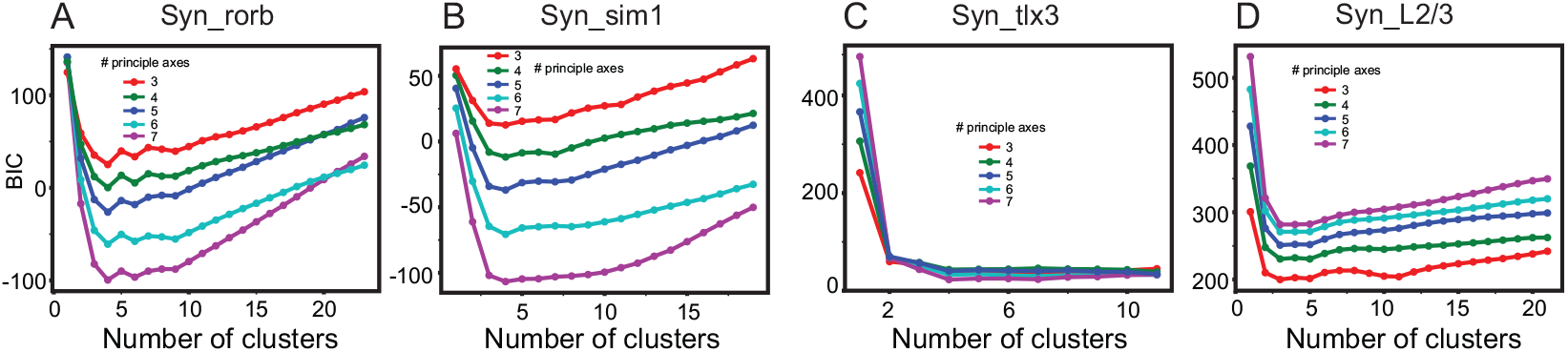
The optimal number of subclasses via x-means clustering. For a range of possible number of subclasses, we conduct k-means clustering 1000 times and calculate the mean values of BIC from them. The different colors represent numbers of principal components selected to reduce the dimensionality of PSP vectors. (A)-(D). The results for four synapse classes (Syn_rorb, Syn_sim1, Syn_tlx3 and Syn_L2/3), respectively.

### 3.2 Fitting synapses to the model

After identifying homogeneous subclasses, we estimate the average PSP vectors from the synapses in subclasses and fit all 12 synapse models (Methods) to them to find the best model of STP for each subclass.

For a synapse subclass, we seek a single synapse model that can account for all patterns of STP observed in multiple stimulation protocols. Thus, the average PSP vectors are independently estimated for all frequencies of presynaptic spikes. Because 8 different pairs of frequency and delay periods are used, there are maximally 8 PSP vectors for each synapse class. However, some synapses are not fully recorded in all 8 conditions, and thus the number of PSP vectors vary over subclasses. Some PSP vectors can be measured incorrectly or affected significantly by noise, and incorporating all PSP vectors will increase the chance of incorrect measurements’ reducing accuracy of STP characterization. To avoid this potential degradation, it is necessary to exclude PSP vectors, which are strongly affected by noise, from STP analysis. As we do not know when the inaccurate measurements occur, we compare PSP amplitudes (i.e., components of PSP vectors) with one another in each stimulation protocol and test whether any change in each PSP vector can be too big to be explained by STP. For instance, if PSPs are measured with artifacts independent of synaptic transmission, their amplitudes are underestimated, inducing abrupt changes from earlier and later PSPs beyond natural changes in STP. To determine whether such abrupt changes exist in PSP vectors, we fit each PSP vector (from subclasses) to the TM model, whose explaining power is well documented. PSP vectors are further analyzed only when *R*^*^2^*^> 0.4. The threshold value is intentionally set to be low to include data for further analysis as much as possible.

After removing PSP vectors which are poorly fit to TM model, we inspect the number of PSP vectors existing in each subclass and select all subclasses including at least two more PSP vectors. 3 subclasses of Syn_L2/3 and 2 subclasses of all other classes satisfy this condition. Then, all 12 models (Methods and Table 1) are fit to the PSP vectors in these subclasses. In this study, we use ‘lmfit’ package [16], the open-source python package developed for nonlinear least-square fit, to find optimal parameters for all 12 models. After fitting, we compare the adjusted *R*^*^2^*^, BIC and Akaike information criterion (AIC) to find the best model for subclasses of synapses (Fig. 2). The best model exists when BIC and AIC are minimized and when the adjusted *R*^*^2^*^ is maximized. For instance, model 5 is the best descriptor for the first subclass of Syn_sim1 (Fig. 2). Fig. 3 shows the best fitting results; one subclass of Syn_L2/3 is not well explained by all 12 synapse models tested and thus not displayed. The parameters from the best models of each subclass are listed in Table 2.

**Table 2:**
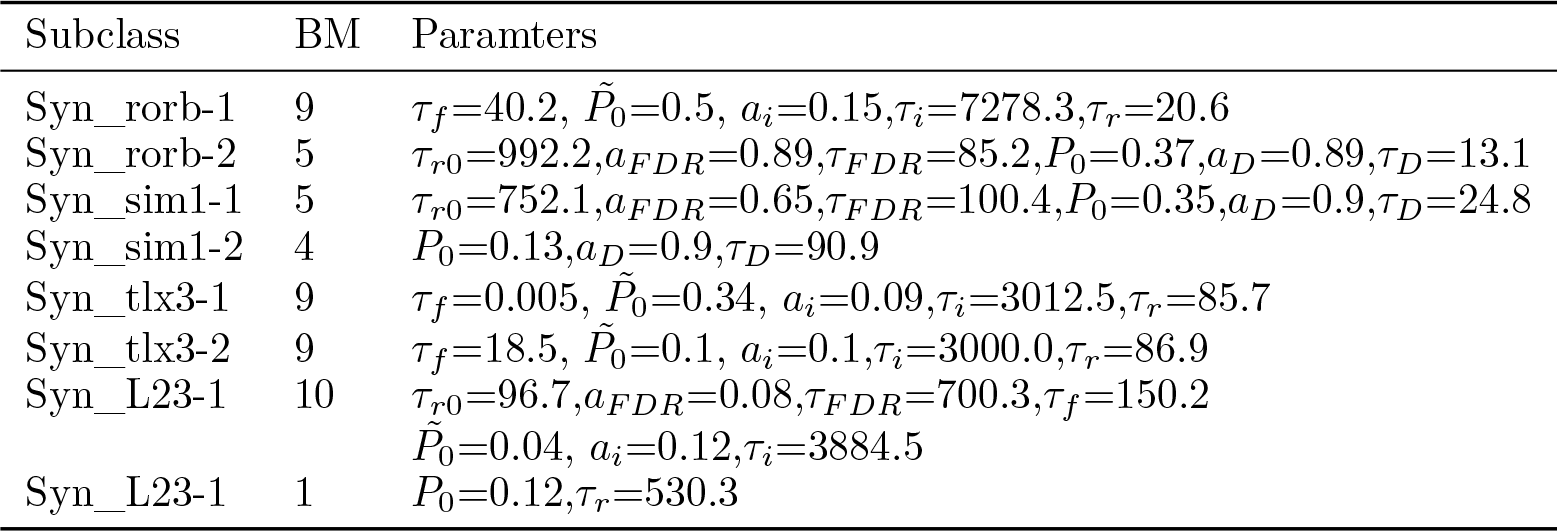
Parameters for each synapse subclass. We list the best model (BM) for each subclass and the estimated parameters for them. The unit of time constants is ms.

**Figure 2:**
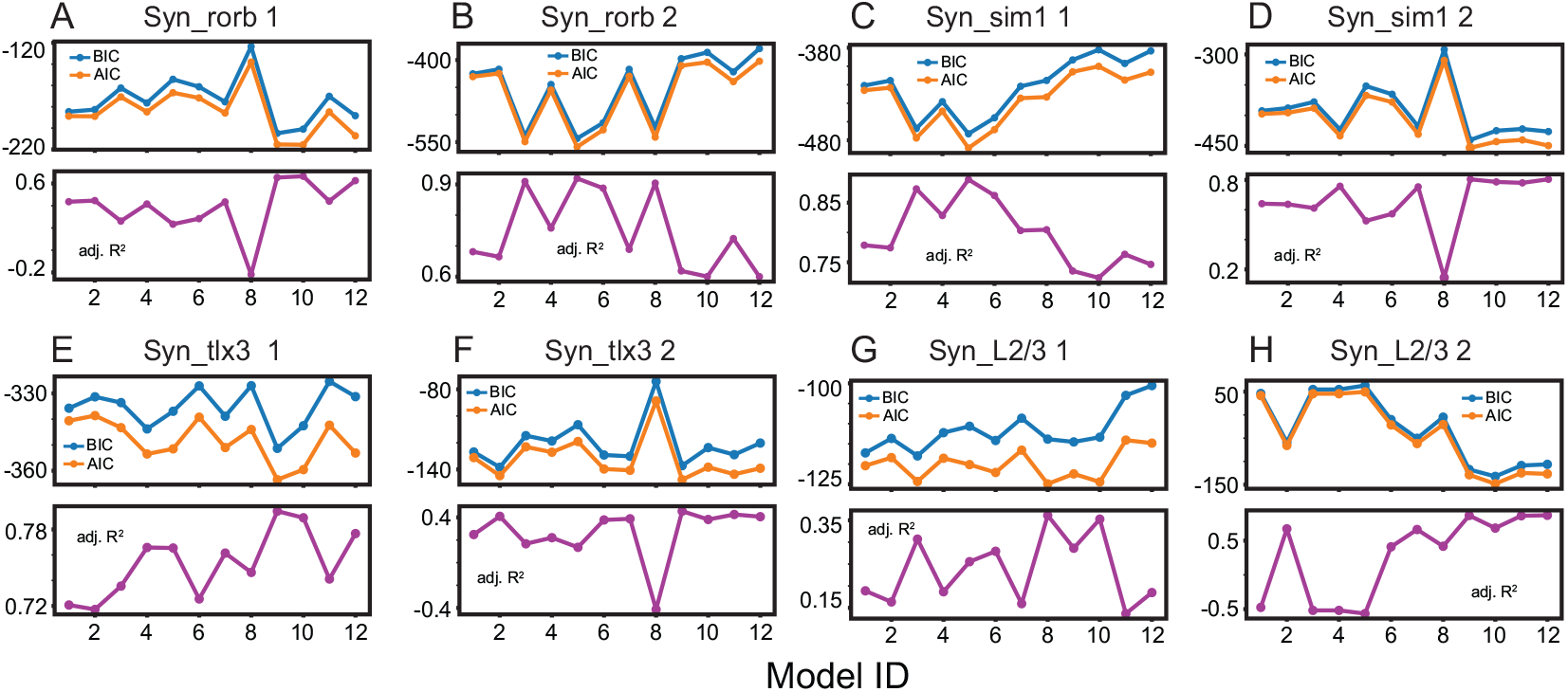
Fitting results with 12 synapse models. In each panel, BIC, AIC and adjusted R2 are illustrated in blue, yellow, magenta, respectively. For each synapse class, two distinct subclasses are displayed in (A)-(H)

**Figure 3:**
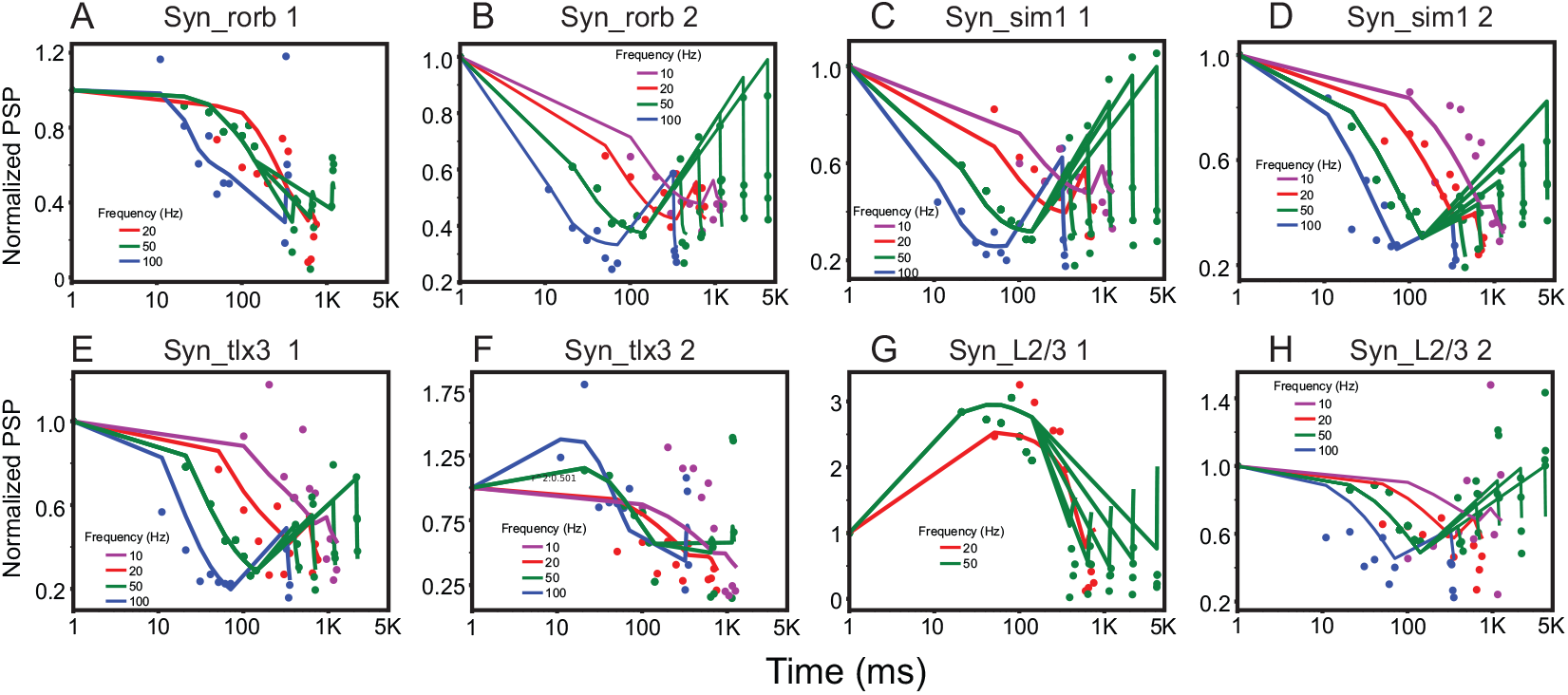
The best fitting results. The fitting results of the best model for each subclass. The dots represent experimental data estimated from subclasses (i.e, PSP vectors), and the lines represent PSP amplitudes predicted with the best synapse models. The color codes are used to show the frequency of presynaptic spikes.

We made a few germane observations. First, each synapse class has at least two subclasses; the number should be considered as an estimate of the lowest bound. Second, the subclasses from the same classes are best described by different models. Only two subclasses of Syn_tlx3 are best described by the same model (model 9). Third, we found two subclasses of Syn_L2/3, while STP of Syn_L2/3 could not be explained by the model in the original study [23]. Further, a subclass of Syn_L2/3 shows a strong facilitation.

### 3.3 Functions of heterogeneous STP

The diversity of STP and heterogeneous subclasses of synapses raise an important question: “Why does the brain have diverse STP?” An earlier theoretical study[1] proposed that depressing synapses allow neurons to be more sensitive to the change in small afferent inputs, which is referred to as ‘gain control’. Since our analyses show diverse patterns of depression and facilitation, we revisit the synapses’ contribution to this gain control by using a network model (Fig. 4A); see supplemental data for details of model implementation. In the model, REC population receives inputs from two populations (*PRJ*_*1*_ and *PRJ*_*2*_). *PRJ*_*1*_ projects rapidly modulating ones at 40 Hz, whereas *PRJ*_*2*_ projects slowly modulating afferent inputs at 10 Hz. The mean values of these modulating inputs are 100 Hz, and the modulation amplitudes are 50 Hz for both rapid and slow modulations. In the model, all neurons are excitatory, and connections with STP are randomly established between neurons at 30% probability. REC neurons also receive 1500 Hz Poisson spikes background inputs via static synapses. The strength of both static and dynamic connections is 100 PA.

**Figure 4:**
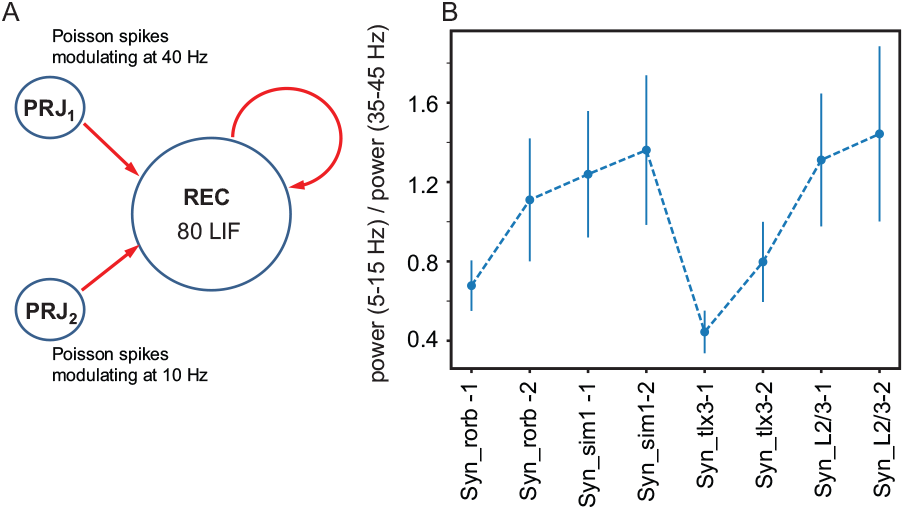
Network simulation results. (A), Model structure. The red arrows represent recurrent synaptic connections with STP in the model. All connections have identical STP estimated for each synapse subclass (Table 2). (B), The ratio of spectral power between 5-15 Hz to the ratio between 35-45 Hz, for each subclass.

If REC neurons respond to 40 Hz modulation, the peak power around 40 Hz would appear in their outputs, and vice versa. Thus, we measure REC neurons’ outputs in the two frequency bands (5-15 and 35-45 Hz), depending on synapse subclasses. For each subclass, we independently instantiate 100 networks and simulate their responses for 2 seconds. In each simulation, we estimate the ratio of spectral power between 5 and 15 Hz to that between 35-45 Hz. That is, the ratios below 1 mean that REC neurons respond more strongly to 40 Hz modulation inputs, and those above 1 mean that REC neurons respond preferably to 10 Hz. Fig. 4B shows the mean values and standard deviations estimated from these 100 simulations depending on subclasses (Fig. 3). For all synapse subclasses, the power ratios are significantly (t-test, *p* < 0.0001) different from 1, suggesting that neurons prefer either 10 or 40 Hz modulation depending on STP.

## 4 Discussion

The dynamics of neural responses in the brain, which are the substrates of our cognitive functions, are controlled by synaptic communications evolving over time. Thus, to better understand the operating principles of the brain, it is imperative to understand STP’s contribution to neural dynamics. It is increasingly clear that a standardized-large-scale survey of synapses is necessary to better understand functions of STP, and the synaptic physiology pipeline of Allen Institute for Brain Science aims to generate such database to fully characterize the properties of synapses including STP with standardized protocols; see [23].

In our study, we develop a workflow to characterize STP using data from Allen Institute for Brain Science synaptic physiology pipeline. Our workflow includes two different steps. First, the clustering of PSP vectors (i.e., the time course of PSP) is employed to identify homogeneous subclasses in the synapse classes. Second, synaptic parameters regarding STP are optimized to account for observed STP patterns. The former is not introduced in the original study [23], and the latter extends their analysis using more comprehensive models. Our new analyses with this newly developed workflow propose the heterogeneity of STP in synapse classes, and we use network models to study potential functions of diverse patterns of STP in the gain control. In the network model, the excitatory neurons’ outputs are modulated by either slowly or rapidly modulating inputs depending on STP, even though neuronal properties are identical for all simulations. This indicates that diverse patterns of STP can tune neurons to respond selectively to inputs at certain frequencies (i.e., rhythms).

### 4.1 Implication of STP in L4

In the model, REC neurons respond strongly to 40 Hz when Syn_rorb-1 (i.e., the first subclass of Syn_rorb) is used to connect neurons, whereas they respond strongly to 10 Hz when Syn_rorb-2 is used, which raises the possibility of heterogeneous functional groups of L4 excitatory neurons. That is, a subset of L4 neurons is tuned to monitor slowly changing inputs, and another set is tuned to rapidly changing inputs. This segregation seems inefficient because many heterogeneous groups would be necessary to fully cover inputs’ temporal ranges. However, it is important to note that this functional segregation may allow L4 neurons to pick up specific inputs more reliably. For instance, when a mouse sees an approaching aerial predator, the predator’s movements may induce visual inputs at certain frequencies. If some L4 neurons of primary visual cortex can be tuned finely to detect such inputs, it will be beneficial for the mouse’s survival, which justifies the cost of having specialized functional groups.

### 4.2 Implications of STP in L5

Although both tlx3-expressing neurons and sim1-expressing neurons are in L5, sim1 neurons project to subcortical areas (Allen Mouse Brain Connectivity Atlas, http://connectivity.brain-map.org/) and tlx3 neurons project to cortical areas [12]. Consistent with this observation, our network simulation results suggest that tlx3 and sim1 neurons (i.e., REC neurons connected with Syn_tlx3 and Syn_sim1, respectively) have distinct response characteristics due to STP. Interestingly, we note that top-down inputs in a beta or alpha frequency band can project onto deep layer neurons [2, 4, 22, 21, 27]. Given the sensitivity of sim1 neurons in the network model to 10 Hz rhythms, we propose that sim1 mediates top-down inputs in low-frequency bands to subcortical areas. Similarly, the sensitivity of tlx3-neurons in the network model raises the possibility that they mediate rapidly changing rhythms that are known to be prominent in bottom-up inputs [7, 5], into other cortical areas.

### 4.3 Future directions

We would like to point out that the current workflow is examined with a limited amount of data. The original study discussed the connectivity in the mouse primary visual cortex, which was estimated from a large set of data on connectivity matrix, but its STP data was much smaller, limiting our testing of the new workflow. Similarly, our network simulation tests a simple scenario, which cannot fully describe the interactions in the actual networks in the brain. For instance, we ignore the interactions between excitatory and inhibitory neurons and the effects of heterogeneity of inhibitory neuron types. However, despite its test with limited STP dataset, we believe that our newly developed workflow and its results can initiate a substantial discussion on standardized analyses of STP, which is in need due to increasing amount of STP experimental data. In the future, we will continuously extend this workflow and network simulations to incorporate new developments reported in relevant literature and in the Allen Institute for Brain Science pipeline. All tools and simulation codes will be available to the public and maintained up-to-date, and a copy of the private repository can be requested without any restrictions.

