## Supplemental Method and Figure for "Functional synapse types via characterization of short-term synaptic plasticity"

### 1 Implementation of network model

To better understand functions of STP, we use a network consisting of leaky integrate-and-fire (LIF) neurons. Inspired by an earlier modeling study [1] proposing that depressing synapses control the gain of neuron responses to afferent inputs, we examine how STP changes neurons' responses to afferent inputs modulating at 40 and 10 Hz, which have been thought to mediate bottom-up and top-down inputs [2, 3, 4, 6, 7], respectively. In doing so, we implement three populations $PRJ_1$, $PRJ_2$ and REC in the network. The first two populations generate outputs varying over time and project them to the third population (REC). The outputs of $PRJ_1$ are rapidly modulating at 40 Hz, whereas the outputs of $PRJ_2$ are slowly modulating at 10 Hz.

We use NEST [5] to implement this network. REC consist of 80 LIF neurons, which are implemented using neuron module 'iaf_psc_exp' in the NEST [5]; all neuron parameters are the same as default values in the module. Both $PRJ_1$ and $PRJ_2$ consist of 20 neurons, each of which generates independent Poisson spikes modulating at 40 Hz or 10 Hz. Specifically, we implement these independent Poisson spike trains by connecting 'parrot' neurons to the 'sinusoidal_poisson_generator' in the NEST [5]. In the NEST, the poisson_generator generates independent spike trains and projects them to target neurons, and parrot neurons relay them to other neurons in the networks.

To generate both slow and fast modulating Poisson spike trains, we use two independent sinusoidal_poisson_generators in the network. The shared parameters of these two devices are 'rate'=100, 'amplitude'=50, but the first generator connected to $PRJ_1$ is set with 'frequency'=40, whereas the second generator connected to $PRJ_2$ is set with 'frequency'=10. During 2-second-long simulations, we measure spikes from all REC neurons and calculate the spectral power of outputs of REC using histogram of spikes from all REC neurons. The resolution of histogram is 1 ms and is smoothed with Gaussian kernel with 4 ms width. STP

Preprint. Under review.

connections are implemented using the extension module of NEST, which is available to the public at the github repository (https://github.com/AllenInstitute/AIBS_synapse).

### 2 Supplemental figure

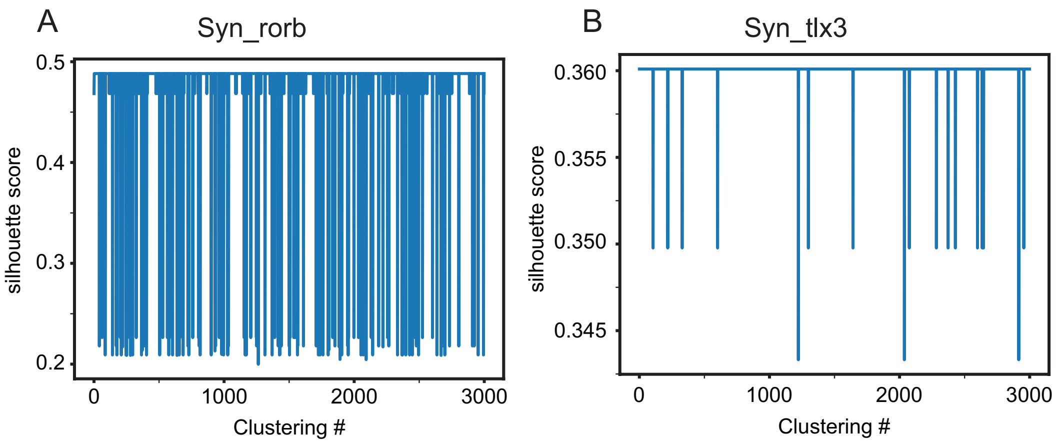


Supplemental Figure 1: Silhouette scores for 3000 clustering with the optimal number of subclasses. The examples of silhouette scores are shown in (A) and (B).

[5] Charl Linssen, Mikkel Elle Lepperød, Jessica Mitchell, Jari Pronold, Jochen Martin Eppler, Chrisitan Keup, Alexander Peyser, Susanne Kunkel, Philipp Weidel, Yannick Nodem, Dennis Terhorst, Rajalekshmi Deepu, Moritz Deger, Jan Hahne, Ankur Sinha, Alberto Antonietti, Maximilian Schmidt, Luciano Paz, Jesús Garrido, Tammo Ippen, Luis Riquelme, Alex Serenko, Tobias Kühn, Itaru Kitayama, Håkon Mørk, Sebastian Spreizer, Jakob Jordan, Jeyashree Krishnan, Mario Senden, Espen Hagen, Alexey Shusharin, Stine Brekke Vennemo, Dimitri Rodarie, Abigail Morrison, Steffen Graber, Jannis Schuecker, Sandra Diaz, Barna Zajzon, and Hans Ekkehard Plesser. NEST 2.16.0. aug 2018.

[6] Anita K Roopun, Fiona E N Lebeau, James Ramell, Mark O Cunningham, Roger D Traub, and Miles A Whittington. Cholinergic neuromodulation controls directed temporal communication in neocortex in vitro. *Frontiers in neural circuits*, 4(March):8, jan 2010.

[7] Xiao-jing Wang. Neurophysiological and Computational Principles of Cortical Rhythms in Cognition. *Physiological Reviews*, pages 1195–1268, 2010.
